# Tripartite networks show that keystone species can multitask

**DOI:** 10.1101/2021.04.01.437523

**Authors:** Sérgio Timóteo, Jörg Albrecht, Beatriz Rumeu, Ana C. Norte, Anna Traveset, Carol M. Frost, Elizabete Marchante, Francisco A. López-Núñez, Guadalupe Peralta, Jane Memmott, Jens M. Olesen, José M. Costa, Luís P. da Silva, Luísa G. Carvalheiro, Marta Correia, Michael Staab, Nico Blüthgen, Nina Farwig, Sandra Hervías-Parejo, Sergei Mironov, Susana Rodríguez-Echeverría, Ruben Heleno

## Abstract

Keystone species are disproportionately important for ecosystem functioning (1,2). However, while all species engage in multiple interaction types with other species, the importance of keystone species is often defined based on a single dimension of their Eltonian niche (3), that is, one type of interaction (e.g., keystone predator). Therefore, it remains unclear whether the importance of keystone species is unidimensional or if it extends across interaction types. We conducted a meta-analysis of tripartite interaction networks to examine whether species importance in one dimension of their niche is mirrored in other niche dimensions, and whether this is associated with interaction outcome, intimacy, or species richness. We show that keystone species importance is positively associated across multiple ecological niche dimensions, independently of species’ abundance, and find no evidence that multidimensionality of keystone species is influenced by the explanatory variables. We propose that the role of keystone species extends across multiple ecological niche dimensions, with important implications for ecosystem resilience and conservation.

**Significance Statement:** Keystone species are often identified by focusing on a single type of interaction (e.g., predation, pollination, herbivory) which contrasts with the multiple roles that species play in biological communities. We conducted a meta-analysis of 18 tripartite interaction networks to explore if keystonness is correlated across the multiple dimensions of species Eltonian niches. Our results suggest that species importance tends to span across multiple interaction types, independently from abundance, which can be key to understand community resilience and collapse in face of multiple threats.

## Introduction

Keystone species play a critical role in ecosystems’ stability (1,2), and are thus a popular concept in ecological research and conservation practice. How these species might affect ecosystem responses to multiple threats is thus a central question in ecology (4). However, advances have been hindered by an excessive compartmentalization of studies focusing on a single function at a time and by the identification of keystone species based on slightly different criteria (reviewed in 5). A frequently used definition characterizes keystone species as those whose proportional importance for a given ecological process (e.g., herbivory, parasitism, pollination, etc.) greatly surpasses that of other more abundant species in the community (2). Due to the intrinsic difficulty of quantifying species contributions to multiple ecological functions at the community level, most research on the role of keystone species has focused on a single ecological function (1,6). However, all species establish different types of interactions with other species in their surroundings, thus playing multiple ecological roles that together define the multiple dimensions of their Eltonian niche (7, 8). For example, many birds include insects and fruits in their diets, thus acting simultaneously as predators and as seed dispersers if the seeds pass unharmed through their digestive tract (9). If insects and fruits are not available, some of these birds might opportunistically consume nectar, eventually acting as pollinators, which reflects another niche dimension (10). In addition, many of these birds will also be hosts to parasites (11) reflecting yet other dimensions of the birds’ Eltonian niche. All these interactions require distinct levels of physiological integration between interaction partners (12, 13), ranging from temporary physical contact between individuals (e.g., predation) to full dependency by one of the partners in at least part of its life cycle (e.g., parasitism). Such a myriad of ecological functions raises the question of whether keystone species are equally important across their multiple niche dimensions or, alternatively, if they tend to be particularly important for a single dimension (and, correspondingly, for a single ecological function).

In order to answer this question, we first need to quantify species roles across multiple niche dimensions under a unified theoretical framework (i.e., a common currency), a task that poses a significant challenge (14). Recent advances in ecological network theory and in the availability of high quality empirical datasets have increased our ability to make sense of the intrinsic complexity of biological communities (15). By simultaneously considering species (nodes) and the interactions (links) that bind them together into functional communities, ecological network analysis has offered a valuable tool to explore the emerging patterns generated by the biotic interactions that define a species’ ecological niche (15). Nevertheless, the study of species ecological interaction networks is still dominated by bipartite networks, where species from two groups are linked by a single type of interaction (16), such as pollination (17), seed dispersal (18), or parasitism (19).

Studies simultaneously exploring species importance across two or more interaction types under a network framework have recently started to emerge (10, 20, 21), but are still rare. Earlier work on the relationships across different types of interactions suggested that important plant species might have greater ratio of mutualistic to antagonistic interactions than expected by chance (16). However, more recent research presents contradictory results, reporting a positive association between the importance of birds as seed dispersers and pollinators (10), but no association in the importance of bats as frugivores and nectarivores (21), or between plants for their mycorrhizal fungi and frugivores (20). Importantly, none of these pairwise comparisons accounted for the effect of species abundance as a key driver of interaction frequency and thus of species importance. Moreover, species have highly variable interaction efficiency across the different interactions in which they participate and vary in their effects upon the fitness of their interaction partners (17, 22).

We now know that both species abundance and richness have positive and independent effects on ecosystem multifunctionality (23, 24), with more abundant species being generally more important contributors to community functioning (25). The effect of abundance emerges from neutral processes, where individuals are expected to contribute equally to community function (26, 27). In turn, species functional importance is the combined result of two components, which ultimately drive interaction effectiveness (28, 29). On the one hand, a quantitative component is mostly affected by species’ abundance and drives the underlying encounter probability between species. On the other hand, a qualitative component stemming from species’ interaction preferences and their morphological, physiological and ecological traits (30–32) and drives species *per-capita* effects. Disentangling these two components is thus essential to correctly identify the role of “keystone species” in their community.

Here, we explore whether species’ functional importance, in terms of their effect on other species, i.e., “keystoneness”, is maintained across different interaction types. This task is not a simple one because the assessment of species effects on each other requires laborious experiments. For that reason, so far only a few remarkable studies have empirically estimated both components of interaction effectiveness at the community level (e.g., 17, 22). Moreover, none of these works did such estimates for more than one niche dimension, thus we still need to rely on frequency-based interaction networks to explore species multifunctionality. We assembled a global dataset of quantitative interaction tripartite networks (Table 1), each composed by two bipartite subnetworks from the same community, coupled by a shared set of species at the interface of the paired networks (Fig. 1A). These networks encompass five distinct interaction types: herbivory, parasitism, seed dispersal, pollination, and mycorrhizal interactions (Table 1). Based on this dataset, we independently quantified the importance of each species coupling two subnetworks, i.e., their importance for each of the two niche dimensions. We did this by estimating species strength (See Materials and Methods), a species-level descriptor calculated as the cumulative sum of each species “dependencies” (the proportion of a species interactions with a given species on the other trophic level), and reflecting each species potential to affect the species in the other trophic level with which it interacts (33). Interaction frequencies are known to overemphasize quantitative over qualitative aspects of interaction effectiveness, though previous work on pollination and seed dispersal systems found that they still provide a suitable proxy of population-level effects of species on their interacting partners (26, 31 but see 29). Yet imperfect, interaction frequencies are a suitable proxy to population-level effects because variations in interaction frequencies due to spatiotemporal fluctuations in species abundances is often larger than fluctuations in *per capita* interaction effects, which are more strongly constrained by species traits (31). In this way, species strength is a suitable indicator of species importance in the community (10, 18) and particularly relevant for being independent from species phylogenetic distances (34).

**Table 1.** Characterization of the 18 datasets included in the meta-analysis. Each dataset is a tripartite network consisting of two bipartite subnetworks (each of them reflecting one dimension of species Eltonian niche), linked by the species at the interface of the subnetworks (more details on each dataset in SI Appendix, Table S1). The types of interaction of each subnetwork were characterized in terms of their interaction outcome (antagonistic vs. mutualistic) and level of intimacy (permanent vs. temporary).

| Data set | Network Levels | Interaction Outcomes | Level of Intimacy | Total species | References |
| --- | --- | --- | --- | --- | --- |
| 01 AZORES | Parasite-Seed disperser-Plant | Antagonistic-Mutualistic | Permanent-Temporary | 63 | (59, Heleno unpublished) |
| 02 AZORES | Herbivore-Plant-Seed disperser | Antagonistic-Mutualistic | Permanent-Temporary | 94 | (60) |
| 03 AZORES | Plant-Herbivore-Parasitoid | Antagonistic-Antagonistic | Permanent-Permanent | 79 | (61) |
| 04 GALAPAGOS | Seed disperser-Plant-Pollinator | Mutualistic-Mutualistic | Temporary-Temporary | 390 | (62, 63) |
| 05 DORSET | Plant-Herbivore-Parasitoid | Antagonistic-Antagonistic | Permanent-Permanent | 51 | (64) |
| 06 NORWOOD | Plant-Herbivore-Parasitoid | Antagonistic-Antagonistic | Permanent-Permanent | 42 | (48, 65) |
| 07 NORWOOD | Herbivore-Plant-Pollinator | Antagonistic-Mutualistic | Permanent-Temporary | 329 | (48) |
| 08 NORWOOD | Plant-Herbivore-Parasitoid | Antagonistic-Antagonistic | Permanent-Permanent | 69 | (48) |
| 09 BORNEO | Plant defender-Herbivore-Plant | Antagonistic-Mutualistic | Permanent-Permanent | 89 | (66) |
| 10 COIMBRA | Parasite-Seed disperser-Plant | Antagonistic-Mutualistic | Permanent-Temporary | 60 | (11, da Silva unpublished) |
| 11 BEIRA | Plant-Galler-Parasitoid | Antagonistic-Antagonistic | Permanent-Permanent | 93 | (67) |
| 12 PURBECK | Parasitoid-Pollinator-Plant | Antagonistic-Mutualistic | Permanent-Temporary | 25 | (19) |
| 13 HAWAII | Plant-Herbivore-Parasitoid | Antagonistic-Antagonistic | Permanent-Permanent | 68 | (68) |
| 14 BRISTOL | Plant-Herbivore-Parasitoid | Antagonistic-Antagonistic | Permanent-Permanent | 690 | (69) |
| 15 POLAND | Seed disperser-Plant-Pollinator | Mutualistic-Mutualistic | Temporary-Temporary | 357 | (49) |
| 16 GALAPAGOS | Parasite-Seed disperser-Plant | Antagonistic-Mutualistic | Permanent-Temporary | 114 | (62, Heleno unpublished) |
| 17 MOZAMBIQUE | Mycorrhiza-Plant-Seed disperser | Mutualistic-Mutualistic | Permanent-Temporary | 107 | (20) |
| 18 NEW ZEALAND | Plant-Herbivore-Parasitoid | Antagonistic-Antagonistic | Permanent-Permanent | 204 | (70, 71) |

**Fig. 1.**
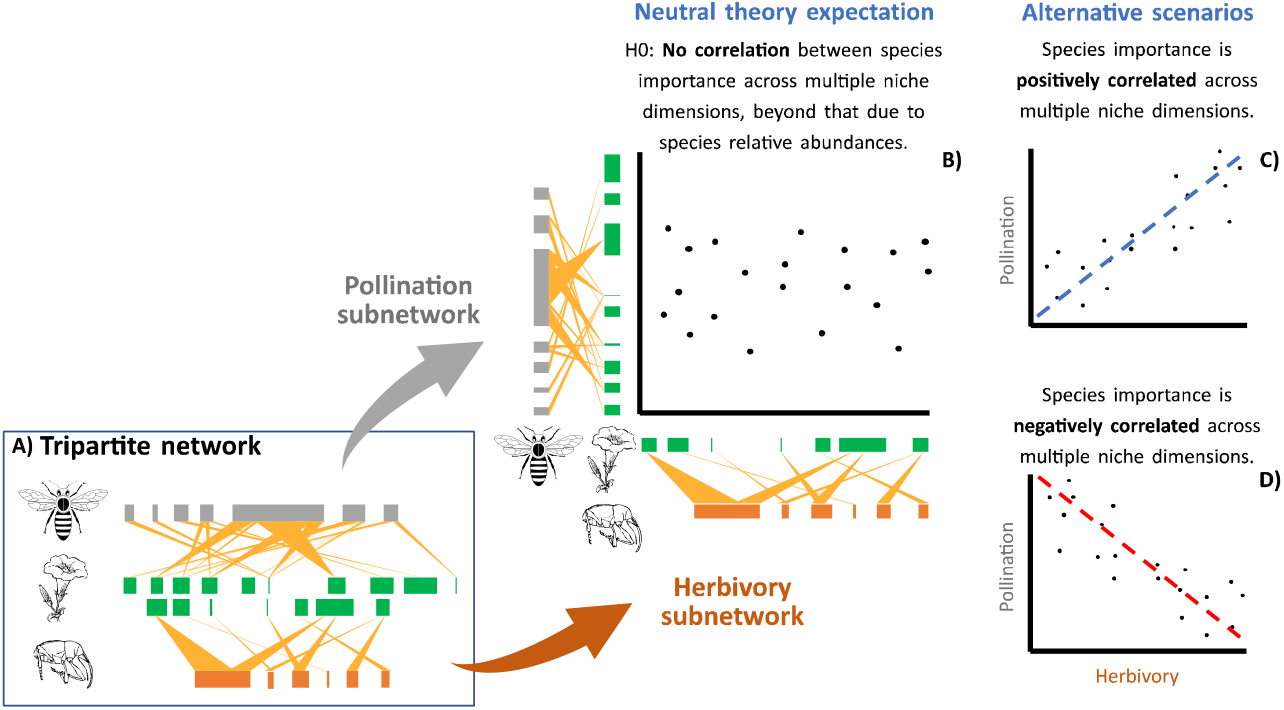
Conceptual model for the importance of species participating in more than one ecological function. Panel A shows a hypothetical tripartite network composed of two bipartite subnetworks: pollination (top) and herbivory (bottom). Grey, green and orange boxes represent species of pollinators, plants, and herbivores, respectively, while yellow lines represent species interactions. Each species importance in the intermediate level (plants in this example) can be assessed by calculating species strength, after normalizing matrices according to the mass-action hypothesis, which accounts for species relative abundances. Panels B, C and D represent alternative hypotheses regarding the correlation of species importance across each paired subnetwork: B) species importance in both subnetworks is independent; C) species importance positively correlated between subnetworks, i.e., keystone species (top right) equally relevant for both dimensions; and D) species importance negatively correlated between subnetworks, i.e., keystone species relevant for either pollination or herbivory. Silhouettes from Open Clipart, under CC0 1.0 licence.

Here, we took a meta-analysis approach to test to what extent a species’ functional importance in one niche dimension is correlated with its importance in a second niche dimension (represented by the interactions in each of the paired bipartite subnetworks, Fig. 1B-D). None of the available tripartite network datasets had available estimates of interaction effectiveness, and very few of the bipartite subnetworks had field-measured abundance data. To circumvent the limitations we followed Staniczenko et al’s approach (30) based on the “mass action hypothesis”, which assumes that interaction frequencies are driven by species abundance (neutrality) and any deviation result from species preferences. This approach allowed us to obtain preference matrices through the estimation of the so-called “effective abundances” by minimizing deviations under the assumption that interaction frequencies were purely driven by mass action (30; See Materials and Methods, and SI Appendix). Firstly, we explored whether removing the effect of species “effective abundances” changes the correlation in species importance between subnetworks. Then, we investigated whether any relationship found was driven by 1) the qualitative outcome of the interaction in terms of its fitness (i.e., whether interactions are mutualistic or antagonistic), 2) the intimacy of the interaction (i.e., whether interactions are temporary or permanent), or 3) community species richness.

## Results

The 18 tripartite species interaction networks had on average 162 species (min. = 25; max. = 690), and 367 links between species (min. = 56; max. = 1414). The tripartite networks had on average 58 species at the intermediate level (min. = 9; max. = 359), of which on average 23 species (min. = 7; max. = 155) participated in both bipartite subnetworks, thus forming the interface between the two subnetworks (SI Appendix, Table S1).

We found a positive and statistically significant overall correlation of the original species strength between paired subnetworks (mean Pearson’s *r* = 0.42, z = 3.34, p < 0.001; Fig. 2, Table 2, and SI Appendix, Table S2 for the correlations of each network). Accounting for species abundances reduced the correlation of species strength between paired subnetworks in 16 of the 18 studies (mean difference = 0.11, paired t-test: *t* = 3.31, df = 17, p = 0.004). However, even after accounting for species abundances the overall correlation of standardized species strength between paired subnetworks remained positive and statistically significant (mean Pearson’s *r* = 0.24, z = 2.42, p = 0.016; Fig. 2, Table 2, and SI Appendix, Table S2 for the correlations of each network).

**Fig. 2.**
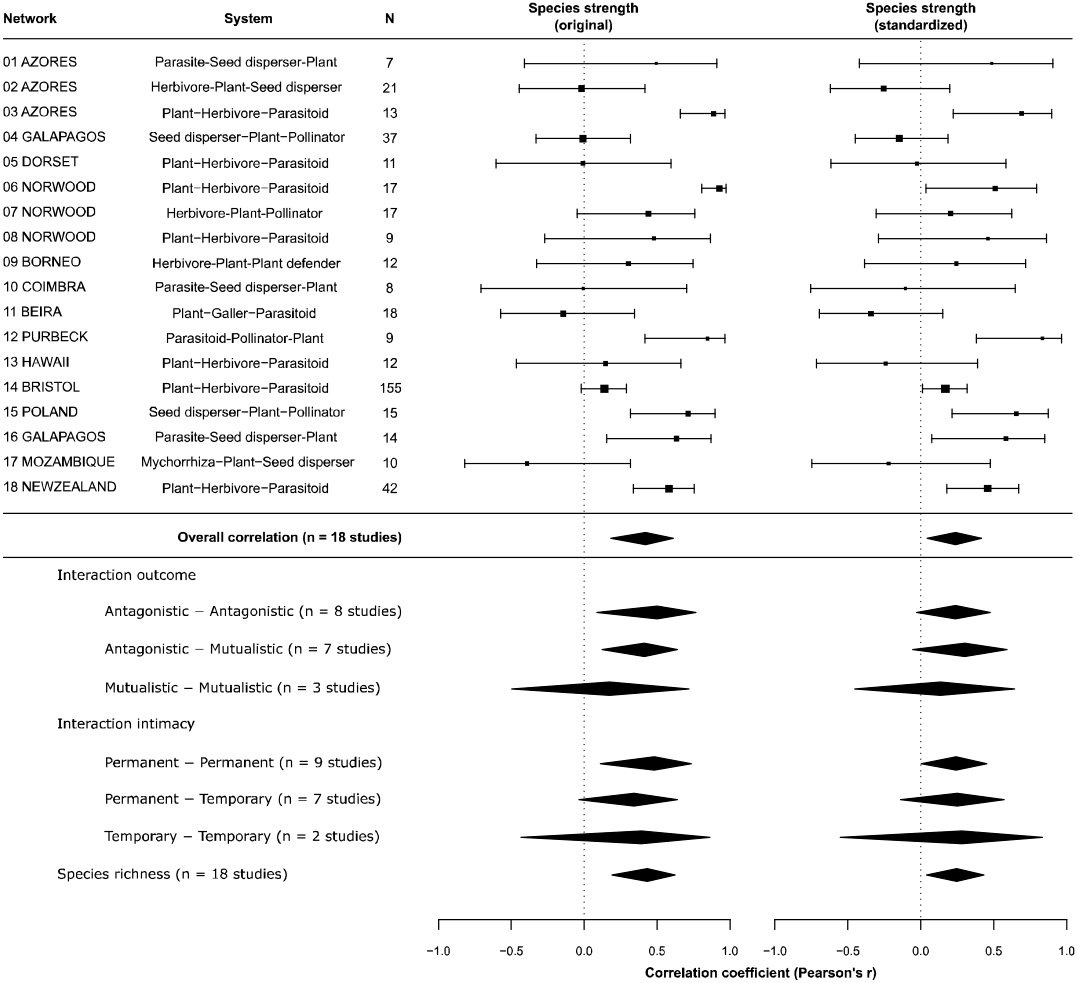
Forest plot with the results of the meta-analysis. Results of the meta-analysis on the Pearson’s correlation coefficient (*r*) of species strength for species participating in paired ecological functions (subnetworks) in 18 tripartite networks (i.e., interface species) (Table 1, and SI Appendix, Table S1). Overall, the importance of species at the interface tends to be positively associated between the two subnetworks, even when the mass action (i.e., abundance) effect is removed (i.e., the standardized species strength), indicating that the keystoneness tends to be maintained across multiple niche dimensions (Fig. 1C). N is the number of species common to both subnetworks. The correlation coefficient of each tripartite network is represented by the square at the centre of the 95% confidence interval and its size is proportional to N. Diamonds represent the overall weighted correlation coefficient and its 95% confidence interval (individual values in SI Appendix, Table S2).

**Table 2.** Results of the meta-analysis of the tripartite networks. The random model is the model without moderators and includes all studies. The subsequent rows present the models for each combination of interaction outcome and intimacy, each with three combinations of interactions, and for species richness. Pearson’s *r* is the overall correlation coefficient [95% confidence interval] of each model; df are the degrees of freedom of the model; *I^2^* is the heterogeneity among correlation coefficients that is not due random sampling variance; n is the number of studies included in each model; *QM* is the amount of heterogeneity accounted by moderators; p is the significance of the Q-test for heterogeneity; and *QE* is the amount of heterogeneity across studies included in the models. For full results of the meta-analytical models see SI Appendix, Table S3. Significant correlations, i.e., those for which the 95% confidence interval did not overlap zero, are given in bold.

|  | Species strength<br>(original) |  | Species strength<br>(standardized by abundance) |  |
| --- | --- | --- | --- | --- |
|  | Pearson's *r* | *I*<sup>2</sup> | Pearson's *r* | *I*<sup>2</sup> |
| Random effects model (no moderators, n = 18) | **0.42 [0.18, 0.61]** | 80.2% | **0.24 [0.05, 0.42]** | 63.2% |
| Test for heterogeneity between studies | QE = 72.403, df = 17, p < 0.001 |  | QE = 39.512, df = 17, p = 0.002 |  |
| Subgroups: Interaction outcome |  |  |  |  |
| Antagonistic + Antagonistic (n = 8) | **0.50 [0.08, 0.76]** | 88.8% | 0.24 [-0.03, 0.48] | 65.1% |
| Antagonistic + Mutualistic (n = 7) | **0.41 [0.12, 0.64]** | 38.9% | 0.30 [-0.05, 0.59] | 53.7% |
| Mutualistic + Mutualistic (n = 3) | 0.17 [-0.50, 0.71] | 81.9% | 0.14 [-0.45, 0.64] | 76.0% |
| Test for difference between moderator subgroups | QM = 0.894, df = 2, p = 0.640 |  | QM = 0.327, df = 2, p = 0.849 |  |
| Subgroups: Interaction intimacy |  |  |  |  |
| Permanent + Permanent (n = 9) | **0.48 [0.11, 0.73]** | 86.6% | **0.24 [0.00, 0.45]** | 58.1% |
| Permanent + Temporary (n = 7) | 0.34 [-0.04, 0.63] | 58.5% | 0.25 [-0.14, 0.57] | 58.6% |
| Temporary + Temporary (n = 2) | 0.39 [-0.44, 0.86] | 86.2% | 0.28 [-0.55, 0.83] | 87.0% |
| Test for difference between moderator subgroups | QM = 0.288, df = 2, p = 0.867 |  | QM = 0.001, df = 2, p = 0.999 |  |
| Species Richness | **0.43 [0.19, 0.62]** | 76.1% | **0.24 [0.05, 0.43]** | 60.3% |
| Test for effect of moderator | QM = 0.621, df = 1, p = 0.431 |  | QM = 0.095, df = 1, p = 0.758 |  |

Nevertheless, despite the positive correlation between species importance across the 18 tripartite networks, we detected a large heterogeneity (See Materials and Methods), indicating that correlations were not consistent across all networks. Heterogeneity in correlation coefficients between studies (*QE*) was higher than expected by chance (original data: *QE* = 72.403, df = 17, p < 0.001; standardized: *QE* = 39.524, df = 17, p = 0.002, Table 2 and SI Appendix, Table S3), and accounted for a relatively high proportion of the total variability between studies (original species strength: *I*^2^ = 80.2%; standardized species strength: *I*^2^ = 63.3%; Table 2 and SI Appendix, Table S3).

The subgroup analyses (See Materials and Methods) indicated that the mean correlations of original and standardized species strength were consistent among subgroups, even if the uncertainty was larger for those combinations of interaction outcome and intimacy that were represented by fewer networks (i.e., mutualistic-mutualistic, n = 3; temporary-temporary, n = 2; Table 2), and being more evident for networks with antagonist or permanent interactions (Fig. 2 and Table 2). Including the effect of community species richness (See Materials and Methods) did not significantly change the overall correlation of species strength between paired subnetworks for neither the original nor the standardized data (mean Pearson’s *r* = 0.43 and 0.24, respectively; Fig. 2, Table 2, and SI Appendix, Table S3). Including the moderators (i.e., interaction outcome, interaction intimacy, and community species richness) accounted for virtually no heterogeneity in the correlations of original and standardized species strength (Table 2 and SI Appendix, Table S3).

## Discussion

Natural communities are bound together by multiple types of biotic interactions. Understanding how species and their interactions couple processes across multiple functional levels is critical to advance our understanding of ecosystem structure and functioning (35), and for predicting the effects of global change on complex ecosystems (36, 37). Our analysis of tripartite networks revealed that species’ overall importance, in terms of their effects on other species, tends to be positively associated across multiple ecological niche dimensions, independently from species abundance. However, by removing the effect of abundance, the strength of this association is attenuated to half of the association found for the original data (Fig. 2, Table 2, and SI Appendix, Table S2). This means that species that are disproportionately important for a particular ecosystem function also tend to play a relevant role for other functions in the community they integrate, thus revealing the multidimensionality of keystone species. This seems to be a general pattern, as we found no evidence that the strength of this positive association depends on community species richness, on whether the interaction is antagonistic or mutualistic, or on whether it is temporary (e.g., pollination or seed dispersal) or permanent (e.g., plant-fungal interactions).

### Species “keystoneness”

It is well established that species abundance is an important driver of their overall effect on ecosystem functioning (23, 24). Under the assumptions of the neutral theory of biodiversity (27), we would expect no significant correlation in species importance across multiple niche dimensions, beyond that explained by species relative abundances. By calculating species strength based on interaction preferences, we were able to estimate and isolate the effect of abundance on interaction frequencies. Ideally, independent field-measured estimates of species abundances should be used to correct frequency matrices in addressing species importance. Unfortunately, these are often not available, as they were not available for most of the datasets used here. Yet the theoretical “effective abundances” proposed by Staniczenko *et al.* (30) showed to be a suitable alternative (SI Appendix, Fig. S1).

In addition to abundance, assessing species “keystoneness” should also consider that neither all species nor all individuals of a species have the same impact on the fitness of their interaction “partners”. For instance, not all pollinators or frugivores are similarly effective in contributing to the pollination or seed dispersal of plants, respectively. A more realistic measure of true interaction effectiveness should then consider the combined effect of quantitative and qualitative components of effectiveness (28, 29, 38). However, our work is limited by the lack of tripartite networks in which pair-wise interactions are based on true estimates of interaction effectiveness. Thus, we had to rely on networks based on interaction frequency, by far the most common type of networks available, as surrogates for population level effect of one species on another. Future studies should incorporate community level estimates of interaction effectiveness to confirm our findings and get closer to the real contribution of species to ecosystem functioning (39).

Despite these general limitations, we show that species abundance contributes to the correlation in species roles across different niche dimensions, as the magnitude of these correlations consistently became weaker after we removed the effect of abundance. However, even after controlling for abundance, we still found a significant positive signal (though half of that found in the original data) of species functional importance across multiple interaction networks (Fig. 2, Table 2, and SI Appendix, Tables S2 and S3). Clarifying the relative importance of all the drivers of species interactions remains a exciting goal in community ecology (7). The results we report here indicate that, beyond species abundances, intrinsic species preferences driven by other non-neutral factors, such as those related to morphological trait-matching (7), physiological constraints (21), temporal and spatial overlap between interacting species (40, 41), and foraging behaviour (42) contribute to species functional importance. Several of these drivers are naturaly associated with species evolutionary and phylogenetic history, such as body size and other morphological traits (7), while others are contingent on ecological (e.g., alternative resources, phenological mismatches, (43)) and behavioural (e.g., fear landscapes (44)) constraints. Contrary to the effect of species abundances (30) there is still no satisfactory method to isolate the effect of phylogenetic relatedness from species preferences on pairwise interactions matrices (45). Nonetheless, species strength is generaly assumed to be largely independent from species phylogeny (34).

### Conservation implications

The identification of keystone species has long been a central conservation priority (1,2,4). Our study provides evidence that the importance of these species is not restricted to single niche dimensions but extends across multiple dimensions of their functional niche. These findings have important implications for conservation planning as they reveal a causal link for coupled responses across distinct – even if apparently disconnected – ecological functions. In particular, the loss of multidimensional keystone species is likely to intensify trophic cascades and rapid community collapse, as these species can trigger parallel responses across different ecological functions, eventually leading to systemic failure (10, 46). On the other hand, the benefits of protecting keystone species will likely extend across multiple ecosystem functions, some of which might not be original conservation targets (47). This suggests that biotic interactions are not only critical to understand specific ecosystem functions, such as predation, disease transmission, or pollination, but also that the dimensionality of species interactions has vital consequences for the structure of entire ecosystems and probably determines their sensitivity to external perturbations and species extinctions (35, 48, 49).

### Concluding remarks

It is important to realize that bipartite networks (those focusing on two groups linked by a single interaction type) represent an abstraction imposed by sampling constraints. Therefore, it is increasingly clear that only by jointly considering the multiple dimensions that characterize species interaction networks we can get closer to understanding the intrinsic complexity of real ecosystems (7, 8, 35). However, very few studies simultaneously quantify multiple interaction types at the same site (e.g., 10, 16, 20). We show that keystone species tend to be disproportionately important across multiple niche dimensions regardless of species abundance, interaction outcome, intimacy, or community species richness. This study represents an important step towards a deeper understanding of the multidimensionality of keystone species. By performing a meta-analysis on a set of networks that include five different interaction types allowed us to escape system-specific conclusions. Our results are constrained by the number and type of empirical tripartite networks available, and by the predominantly incomplete assessment of interaction effectiveness in community-level studies, that still prevent us from getting closer to true measures of effective species “keystoneness”. The rapid advances on the compilation of more, larger, and more detailed datasets, incorporating multiple ecosystem processes under a unified network framework will likely foster a deeper understanding of how keystone species shape ecosystem structure, function, and resilience.

## Methods

### Data set

We assembled a global dataset comprising 18 quantitative tripartite networks, each composed of two bipartite subnetworks (i.e., with two levels that interact with each other), representing two dimensions of species Eltonian niches. Overall, these networks encompass five distinct interaction types (i.e., dimensions): herbivory, parasitism, seed dispersal, pollination, and mycorrhiza (Table 1 and SI Appendix, Table S1). Sampling of both subnetworks coincided in time and space for each tripartite network. If studies included data from nearby plots/sites, these were pooled together, after checking if such pooling made biological/ecological sense (i.e., if those species and interactions can be considered as part of the same biological community).

To explore the potential underlying mechanisms explaining eventual correlations between species importance across their niche dimensions, we characterized each tripartite network regarding the interaction outcome and intimacy of the interactions on each subnetwork, as well as community species richness (Table 1), using these as moderators (i.e., variables driving the variation between studies) in a metaanalysis. Interaction outcome was classified as either antagonistic or mutualistic, resulting in three combinations of outcomes in the tripartite networks: antagonistic-antagonistic, antagonistic-mutualistic, and mutualistic-mutualistic (Table 1). Interaction intimacy describes “the degree of physical proximity or integration of partner taxa during their life cycles” (12). Due to the limited number of tripartite networks available, we followed a conservative approach and classified the degree of interaction intimacy as permanent (high intimacy) or temporary (low intimacy) (13, 35). Permanent interactions are those where one of the partners is physically or physiologically dependent of the other for a significant proportion of their life cycles (e.g., the interaction between mycorrhizal fungi and plants, or parasitoids and their hosts), and temporary interactions are those where such dependencies are restricted to short periods of phenological matching (e.g., the interaction between plants and their pollinators or seed dispersers). This classification resulted in three combinations of intimacy levels: permanent-permanent, permanent-temporary, and temporary-temporary. We defined community species richness as the total number of species that have been recorded across the two subnetworks of each tripartite network (Table 1).

### Estimating species importance

To quantify the importance of each species for each dimension of their niche (i.e., the two subnetworks), we focused on species at the interface of the two subnetworks and calculated their species strength (33). This metric is a particularly useful species-level descriptor that quantifies the cumulative importance of species to the entire assemblage of interacting partners in the other trophic level (10, 18, 26, 50, 51). For example, in an herbivore-plant-pollinator network, species strength was computed twice for each plant: firstly, reflecting its importance as a resource for the pollinator community, and secondly, reflecting its importance as a resource for the herbivore community. For each subnetwork, we constructed an interaction matrix **A** (*I* × *J*), in which each cell *a_i,j_* describes the frequency of interactions between *I* species at the network interface (e.g., plants) and *J* species in the other network level (e.g., pollinators). Let 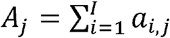 the total number of interactions of species *j*. The dependency of species *j* on species *i* is then denoted as 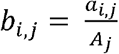. The species strength of species *i* at the network interface is defined as the sum of the dependencies of the *J* species on species *i*: 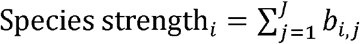. Species strength estimated from the empirical raw data is hereafter referred to as original species strength, and was calculated in R 4.0.2 (52) with package *bipartite v2.08* (53).

### Standardizing interaction matrices and species preferences

To understand whether accounting for species abundances would change the association of species importance between subnetworks, we then standardized all species interaction matrices according to the “mass action hypothesis” (following 35). This hypothesis assumes that species abundances drive the probability of encounter between two species, and that deviations from this assumption are due to species interaction preferences (30). This standardization allowed us to recalculate species importance based on a new set of “interaction preference matrices” that excluded the effect of estimated abundances (named “effective abundances” in (30) on interaction frequencies. As the networks included in this study were assembled by different researchers, with different initial goals and using different sampling methods, this standardization has the additional advantage of converting all interaction frequencies into the same currency.

For a quantitative species interaction matrix **A**, each entry *a_i,j_* of this matrix encodes the weight of the interaction between species *i* and *j* that can be decomposed into: species interaction preferences (*γ_i,j_*) and a mass action effect (*X_i_X_j_*, i.e., the product of the “effective abundances” of the interacting species). Thus, *a_i,j_* = *γ_i,j_X_j_X_j_*, from where interaction preference matrices (i.e., that exclude the effect of species “effective abundances”) can be derived (SI Appendix). It is worth mentioning that “effective abundances” are not necessarily equivalent to real field-measured abundances. They are best-fit estimations that provide the closest approximation of data to the mass action hypothesis, thus representing the theoretical abundance of the interacting species (30). To understand to what extent estimated “effective abundances” reflect measured abundances, we calculated Pearson’s correlation coefficients between these two variables, whenever independent measures of abundance were available for the species at the interface between two bipartite subnetworks (n=7 bipartite subnetworks). The results show that for most datasets, there is a moderate to strong positive correlation between measured abundance and “effective abundance” (r = 0.56 ± 0.20 (mean ± s.d.); SI Appendix, Fig. S1).

Interaction preference matrices were calculated using an adaptation of the R function *GetPreference* from (30). If interactions are exclusively driven by mass action effects, and no deviations occur, the preference matrix would be binary. The existence of interaction preferences is indicated by *γ_i,j_* > 1, an averted interaction is indicated by *γ_i,j_* < 1, and *γ_i,j_* = 1 reflects no deviations from the mass action effect. Interaction preferences *γ_i,j_* thus represent the *per-capita* interaction strength between species *i* and *j*. We then recalculated species strength based on the matrices standardized by “effective abundances”, hereafter referred to as standardized species strength.

### Meta-analysis

For those species at the interface of each tripartite network (i.e., intermediate level), we calculated the Pearson’s correlation coefficient between their values of species strength for the two subnetworks, using both the original and standardized species strength matrices. Species strength often presents a skewed distribution, and therefore was log-transformed. We used a paired t-test to assess if the correlations based on the original and standardized measures of species strength differed systematically.

To estimate the overall correlation across all tripartite networks we took a meta-analysis approach and estimated the associated variance and 95% confidence intervals. This method also weights each effect size (here, Pearson’s correlation coefficients) according to its sample size and variance, and is more robust against type II errors, i.e., falsely rejecting the presence of true effects These advantages offered by meta-analysis, compared to other statistical methods (e.g., hierarchical models), make it more appropriate to use on aggregated effect sizes coming from independent studies (here, each network) with heterogeneous variances (54). Pearson’s coefficients were standardized using a Fisher’s r-to-Z transformation (SI Appendix), which stabilizes the variance of the coefficients.

First, we implemented a random-effect model without moderators, which assumes that effect sizes come from different populations (55). In this model the contribution of each study is weighted by its estimated sampling variance, calculated based on the sample size of each study (i.e., the number of species common to both subnetworks of each tripartite network), thus giving higher influence to studies with larger sample sizes over those with lower sample size, which for that reason may produce less precise coefficients (54, 56) (SI Appendix). The transformed effect sizes (i.e., Z-transformed Pearson’s coefficients) are then used to calculate an average effect size, with each correlation coefficient weighted by the inverse of the within-study variance of the study from which it came (SI Appendix). This results in individual z-values with small variances having greater weights than those with large variances (56). A correlation was considered statistically significant whenever the 95% confidence interval of the correlation coefficients did not overlap zero. In reporting results and their visual representation, Pearson’s coefficients and corresponding confidence intervals were back-transformed (SI Appendix).

Second, we conducted a subgroup analysis to estimate the pooled correlation coefficient for each combination of interaction outcome (i.e., antagonistic-antagonistic, antagonistic-mutualistic, mutualistic-mutualistic), and for each combination of interaction intimacy (i.e., permanent-permanent, permanent-temporary, temporary-temporary). Finally, we included interaction outcome and intimacy as factorial moderators into a mixed effects model to test whether magnitude and sign of the correlation differed between the different combinations of interaction outcome or intimacy. We also added species richness as a continuous moderator in a mixed effect model to test its effect on the correlation coefficients. To avoid model overfitting, we included each moderator at a time. We used the Cochran’s Q-test (57), to test whether correlation coefficients were heterogeneous across the tripartite networks, with a significant result indicating the presence of heterogeneity (*QE*), i.e. the existence of differences in the magnitude and sign of the correlation coefficients between subgroups of studies. We estimated the *I^2^* statistic to quantify the proportion of the total variance resulting from true heterogeneity among studies, i.e. differences between tripartite networks not resulting simply from random sampling variance (57). In models with moderators, the Q-test tests for the presence of significant heterogeneity accounted for by the different levels of the moderator variables (*QM*), i.e., differences in correlation coefficients between the different combinations of interaction outcome and intimacy. Finally, we estimated *R^2^* to quantify the proportion of heterogeneity explained by these moderators. All meta-analysis procedures were conducted using the R package *metafor v2.1-0* (58).

## Acknowledgments

ANC was supported by the Portuguese Science Foundation – FCT/MCTES (transitory norm contract DL57/2016/CP1370/CT89). AT was supported by the Spanish Ministry of Science; Innovation and Universities (PID2020-114324GB-C21). BR was supported by the ‘Juan de la Cierva Incorporación’ fellowship from the Spanish Ministry of Science; Innovation and Universities (IJCI-2017-33475) and the University of Cádiz (UCA/REC17VPCT/2021). SHP was supported by the Spanish Ministry of Economy, Industry and Competitiveness (CGL2017-88122-P). EM, FALN, MC, RH, SRE and ST were supported by the Portuguese Science Foundation – FCT/MCTES (grants PTDC/AAG-REC/4896/2014, SFRH/BD/130942/2017, CEECIND/00092/2017, SFRH/BD/96050/2013, IF/00462/2013, CEECIND/00135/2017, respectively, and all by grant UID/BIA/04004/2020). LPS was supported by the Portuguese Science Foundation – FCT/MTCES (CEECIND/02064/2017). LGC was supported by the Portuguese Science Foundation – FCT/MTCES via the EU Lisbon 2020 program (project EUCLIPO-028360), and by the Brazilian National Council for Scientific and Technological Development (CNPq. Universal 421668/2018-0; PQ 305157/20183).

## Supplementary Information for

### Appendix S1

#### Matrix standardization and species preferences

In order to make networks comparable and to estimate interaction preferences that are independent of species relative abundances we standardized all networks according to the mass action hypothesis (following 1). This also has the advantage of having all interaction frequencies under the same currency. The mass action hypothesis states that the species local abundances drive the probability of encounter between two species, and that deviations from this are due to species interaction preferences (1).

If we have a quantitative species interaction matrix **A**, each entry *a_i,j_* of this matrix encodes the strength of the interaction between species *i* and *j* that can be decomposed into: species interaction preferences *(γ_i,j_*) and mass action (*X_i,X_j__*, the product of the effective abundances of the interacting species), that is

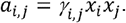

We can then remove the effect of species effective abundances and rescale matrices to obtain estimates of species preferences

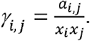

If an interaction is not recorded *a_i,j_* = 0 then *γ_i,j_* = 0, otherwise, when an interaction is present, *a_i,j_* > 0 and *γ_i,j_* ≠ 0. In the latter case, we apply a logarithm transformation obtaining

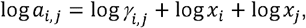

which under the mass action hypothesis is log *γ_i,j_* = 0, thus *γ_i,j_* = 1.

We can re-arrange the log-equation as follows

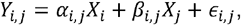

where *α_i,j_* and *β_i,j_* are regression coefficients, *X_i_* and *X_j_* are estimates of the abundances of the interacting species and are unknown, and *ε*_i,j_ is an error term. We seek to minimize the errors *ε* in order to find the best-fit for the estimates of *X_i_* and *Y_j_*, The *γ*-term represent the errors (*ε_i,j_*), and its estimation is constrained to be as close as possible to 1, because log *γ_i,j_* are minimized during regression. In case no errors exist, the standardized matrix is binary with recorded interactions having *γ_i,j_* = 1, and interactions are neutral and result exclusively from the probabilistic encounter between species, i.e. mass action effect. If errors *ε* are present, the solution for the best-fit for *X*_i_ and *X_j_* will result in at least some interaction having *γ_i,j_* ≠ 0, with interaction preferences being indicated by *γ_i,j_* > 1 and less preferred interactions indicated by *γ_i,j_* < 1.

#### Fisher’s r-to-z transformation

Pearson’s coefficients were standardized using a Fisher’s r-to-Z transformation, which stabilizes the variance of the coefficients (2), and obtained through

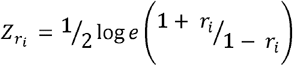

*r_i_* is the Pearson correlation coefficient of the individual studies, which has an approximate normal distribution with variance

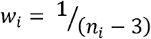

*n_i_* is sample size of the individual studies (in the present study the number of species participating in both functions and from which the correlations where calculated).

#### Average effect size

Transformed effect sizes (i.e., transformed Pearson’s correlation coefficients) are then used to calculate an average effect size, with each correlation weighted by the inverse of the within-study variance (*w_i_*) of the study from which it came from.

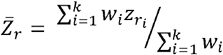

Thus, giving individual z-values with small variances greater weights than those with large variances (3).

#### Back-transformation to Pearson’s correlation coefficient

Pearson’s coefficients and corresponding 95% confidence intervals were back-transformed according to

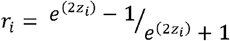

---

**FIG. S1.**
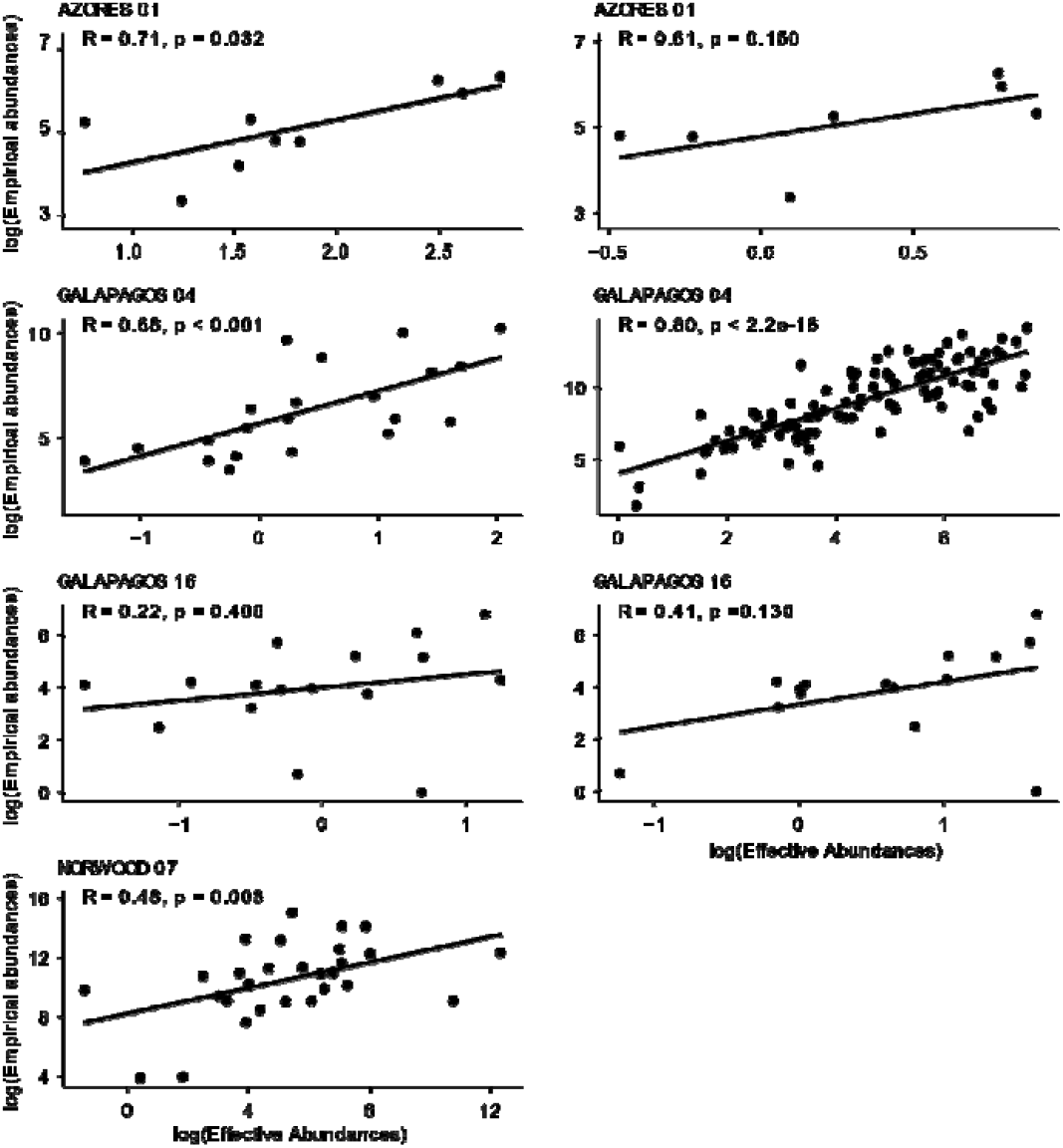
Correlation between the abundance of interaction partners’ resource and effective abundances. Abundance values were log-transformed (see Matrices standardization and species preferences in Methods section and Appendix S1 above). AZORES 01: abundance of birds for parasites (left) and for dispersed plants (right); GALAPAGOS 04: abundance of fruits for seed dispersers (left) and flowers for pollinators (right); GALAPAGOS 16: abundance of birds for parasites (left) and for dispersed plants (right); NORWOOD 07: abundance of plants for herbivores.

**Table S1.** Location and geographic coordinates, network levels of interacting species, total number of species, number of species at the intermediate level (intermediate species), number of common species at the interface of the bipartite subnetworks (interface species), number of links, and size of each bipartite subnetwork of the tripartite networks used in the meta-analysis.

| Data set | Location | Latitude/Longitude | Network levels | Total species | Intermediate species | Interface species | Links | Subnetwork size |
| --- | --- | --- | --- | --- | --- | --- | --- | --- |
| 01 AZORES | Azores, Portugal | 37.794/-25.191 | Parasite-Seed disperser-Plant | 63 | 9 | 7 | 87 | 9x13-7x41 |
| 02 AZORES | Azores, Portugal | 37.794/-25.191 | Herbivore-Plant-Seed disperser | 94 | 53 | 21 | 165 | 31x36-41x7 |
| 03 AZORES | Azores, Portugal | 37.794/-25.191 | Plant-Herbivore-Parasitoid | 79 | 37 | 13 | 110 | 31x36-13x12 |
| 04 GALAPAGOS | Galapagos, Ecuador | -0.738/-89.882 | Seed disperser-Plant-Pollinator | 390 | 163 | 37 | 1414 | 84x21-110x212 |
| 05 DORSET | Dorset, UK | 50.823/-1.830 | Plant-Herbivore-Parasitoid | 51 | 25 | 11 | 87 | 25x11-11x15 |
| 06 NORWOOD | Bristol, UK | 51.313/-2.321 | Plant-Herbivore-Parasitoid | 42 | 19 | 17 | 91 | 19x6-16x16 |
| 07 NORWOOD | Bristol, UK | 51.313/-2.321 | Herbivore-Plant-Pollinator | 329 | 77 | 17 | 540 | 30x28-47x241 |
| 08 NORWOOD | Bristol, UK | 51.313/-2.321 | Plant-Herbivore-Parasitoid | 69 | 28 | 9 | 60 | 28x30-9x11 |
| 09 BORNEO | Danum Valley, Borneo | 4.967/117.800 | Plant defender-Herbivore-Plant | 89 | 13 | 12 | 180 | 13x43-13x32 |
| 10 COIMBRA | Coimbra, Portugal | 38.950/-9.300 | Parasite-Seed disperser-Plant | 60 | 39 | 8 | 158 | 18x6-15x29 |
| 11 BEIRA | Beira Litoral, Portugal | 40.221/-8.890 | Plant-Galler-Parasitoid | 93 | 31 | 18 | 56 | 31x22-18x40 |
| 12 PURBECK | Ilse of Purbeck, UK | 50.616/-2.043 | Parasitoid-Pollinator-Plant | 25 | 9 | 9 | 80 | 9x12-9x4 |
| 13 HAWAII | Hawaii, USA | 23.417/-159.617 | Plant-Herbivore-Parasitoid | 68 | 29 | 12 | 94 | 28x32-13x7 |
| 14 BRISTOL | Bristol, UK | 51.479/-1.984 | Plant-Herbivore-Parasitoid | 690 | 359 | 154 | 1312 | 357x139-157x192 |
| 15 POLAND | Białowieża, Poland | 52.824/24.007 | Seed disperser-Plant-Pollinator | 357 | 15 | 15 | 1220 | 15x34-15x308 |
| 16 GALAPAGOS | Galapagos, Ecuador | -0.738/-89.882 | Parasite-Seed disperser-Plant | 114 | 36 | 14 | 241 | 15x13-16x84 |
| 17 MOZAMBIQUE | Gorongosa, Mozambique | -18.943/34.373 | Mycorrhiza-Plant-Seed disperser | 107 | 19 | 10 | 202 | 16x69-16x16 |
| 18 NEW ZEALAND | Nelson-Marlborough, New Zealand | -41.484/173.006 | Plant-Herbivore-Parasitoid | 204 | 90 | 42 | 502 | 68x76-42x61 |

**Table S2.**
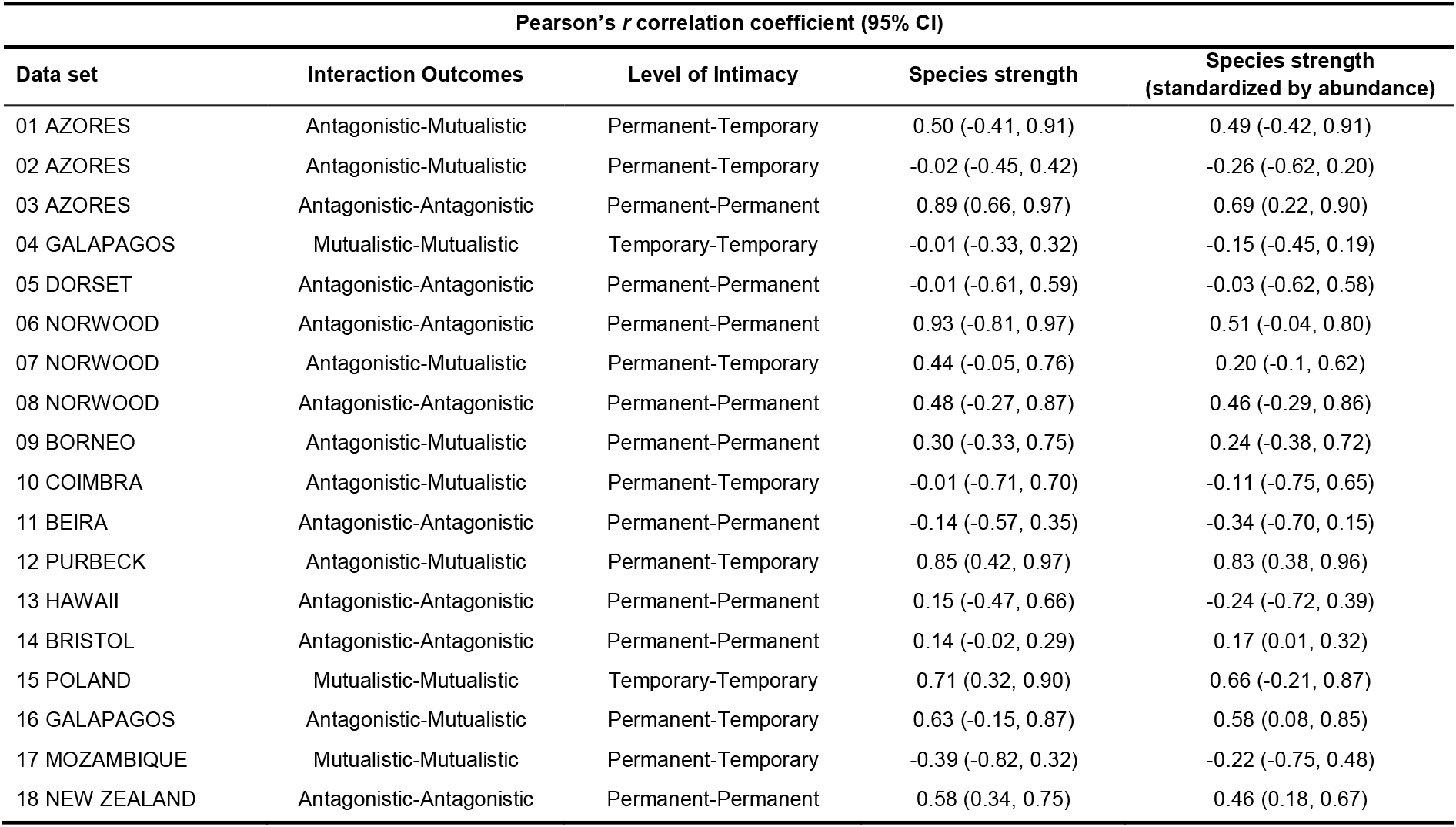
Pearson’s *r* correlation coefficients, with the 95% confidence interval in brackets for each tripartite network included in the meta-analysis of unstandardized and standardized species strength for species participating in both ecological functions.

**Table S3.**
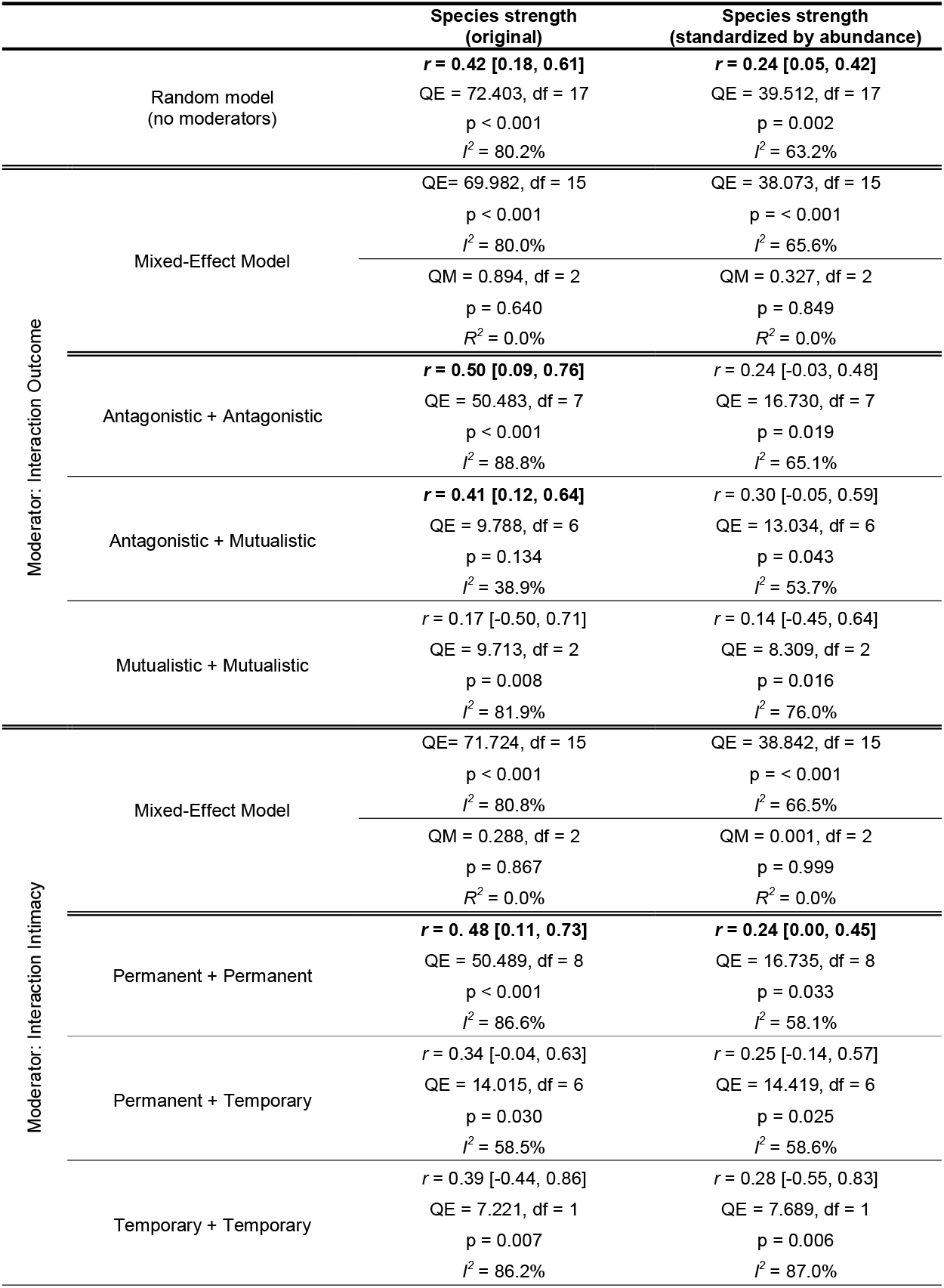
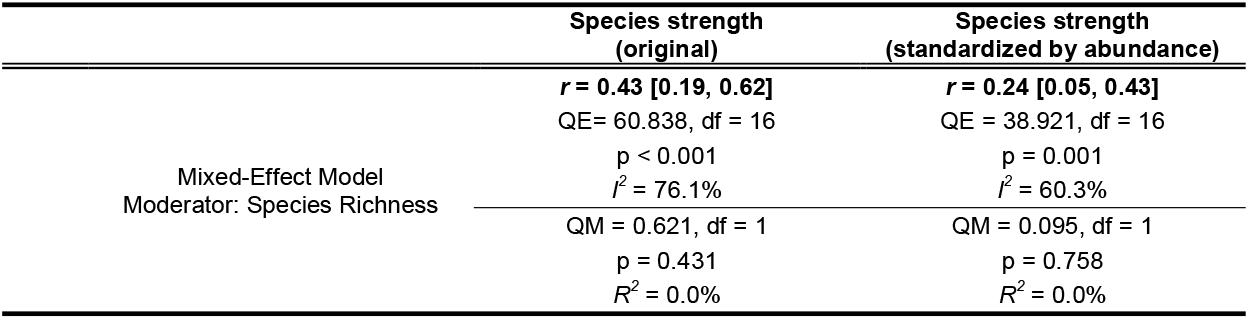
Meta-analysis full results: *r* – Pearson’s correlation coefficient [95% confidence interval] of each model; *QE* – heterogeneity across studies; *I^2^* – % heterogeneity among correlation coefficients not due to random sampling variance; *QM* – heterogeneity accounted by moderators; df – degrees of freedom; p – significance of the Q-test for heterogeneity; *R^2^* – % heterogeneity accounted for by moderators (each with three levels). Correlations for which the 95% confidence interval did not overlap zero are given in bold.

